# The architecture of amyloid fibrils formed by a human tau-derived hexapeptide VQIVYK

**DOI:** 10.1101/2025.03.11.642642

**Authors:** Irene del Mar Farinas Lucas, Youssra K. Al-Hilaly, Liisa Lutter, Wei-Feng Xue, Louise C. Serpell

## Abstract

The sequence _306_VQIVYK_311_ is an aggregation prone region of the tau protein implicated in driving assembly of tau into paired helical filaments. These filaments accumulate as intraneuronal neurofibrillary tangles in Alzheimer’s disease and a range of tauopathies. Here, we demonstrate that VQIVYK forms highly ordered fibrillar samples after prolonged incubation at room temperature. Remarkably, aligned fibre bundles give rise to unusually detailed and highly oriented X-ray fibre diffraction patterns. Analysis of these patterns provide a model of the core protofilament structure that satisfies the experimental diffraction data. This structural model is consistent with data from X-ray crystallography of microcrystals and conforms to the cross-beta architecture that defines amyloid, but diffraction data analysis shows a highly twisted filamentous protofilament architecture. Analysis of individual fibril envelopes by 3D contact point reconstruction atomic force microscopy reveals a diverse polymorphous population with a major fibril morphology of apparent cylindrical fibrils, and morphological subpopulations of fibrils with clear left-hand twisting patterns, despite the commonality of the core structure indicated by the detailed diffraction pattern. Together, these data suggest that VQIVYK amyloid fibrils form a polymorphous amyloid population by assembly of highly ordered protofilaments and provides molecular information regarding amyloid fibril twist.

## Introduction

Amyloid fibrils are defined by the organisation of β-strands that run perpendicular to the fibre axis and are hydrogen bonded along the length of the fibre axis to generate a highly ordered architecture [1]. The β-sheets are associated via interdigitation of the amino side chains to result in a molecular structure with high stability and impressive tensile strength [2-4]. Amyloid fibrils are well known for their association with protein misfolding diseases [5, 6] which include neurodegenerative diseases like Alzheimer’s disease [7] and systemic amyloidoses such as Senile Systemic Amyloidosis [8]. Functional amyloid fibrils provide adhesion and protection in many organisms such as curli in certain bacteria [9] and PMel17 in human melanosomes [10] as well as playing an important role in controlling memory [11] and in viral infection [12]. Due to their inherent strength, amyloid fibrils have been developed as functional materials in a number of bionanomaterial applications [13-15].

Tau is an amyloidogenic protein that self-assembles to form paired helical filaments (PHFs) that are found intracellularly in the neurons of Alzheimer’s brain tissue [16]. PHF are amyloid structures that display all the defining characteristics of a cross-β structure [17]. Full length tau is mostly an intrinsically disordered protein which exists natively unfolded, but several regions of tau have been identified that may serve to drive self-assembly to form fibrils. These are hexapeptide sequences which have been named PHF6 and PHF6* and have the sequences VQIVYK and VQIINK [18-20]. The crystal structures of these peptides derived from fibrillar microcrystals show the classical cross-β structure [19] with a steric zipper architecture similar to fibrous crystals formed by other short amyloidogenic peptides [3, 19]. More recently, cryo-electron microscopy (cryo-EM) has provided a 3D structure for the tau paired helical filament, revealing the core region of the filaments extending from tau 304 to 380 with a parallel β-sheet organisation, whereby each peptide is stacked above an identical sequence [21] The hexapeptide _306_VQIVYK_311_ is found in a β-strand that pairs across the sheet and interdigitates with a peptide _378_FTLKHT_373_ across an antiparallel zipper in *ex vivo* fibril structures extracted from tissue of patients with both Alzheimer’s disease and chronic traumatic encephalopathy[22, 23]. In *ex vivo* structures from corticobasal degeneration brain tissue, VQIVYK forms part of the central four-layered cross-β packing [24] but in all tauopathy filament structures, the sequence is arranged in a parallel β-sheet. These structures point to an important core role of the VQIVYK region and understanding its propensity to form amyloid fibrils alone may provide insight into the parallel stacking arrangement of a range of disease-relevant tau fibril structures and their potential for polymorphic assembly.

Here, we have investigated the filament structure formed from the self-assembly of the VQIVYK peptide, which is not resolved to date. VQIVYK has been widely observed to form amyloid fibrils spontaneously in aqueous buffer and water [25]. We report that following a prolonged incubation, VQIVYK forms exceptionally well-ordered amyloid fibril cores as indicated by an unusually detailed X-ray fibre diffraction pattern that is not typically seen for amyloid fibrils. AFM reveals a polymorphic population, dominated by a smooth filament morphology without detectable twist, and subpopulations with left-handed twisted fibrils. X-ray fibre diffraction patterns were found to provide a very detailed diffraction fingerprint which was analysed alongside complementary atomic force microscopy imaging data with individual filament structural analysis by 3D contact point reconstruction (CPR-AFM). The integrated approach generated a model filament structure that describe both the commonality and the individuality of VQIVYK fibrils and suggest lateral protofilament assembly as a source of structural polymorphism.

## Results and discussion

### Prolonged incubation of VQIVYK results in long straight and twisted fibril polymorphs

Negative stain TEM of fibrils formed from VQIVYK revealed long, straight fibrils with a smooth appearance after very prolonged incubation (approximately 2 years). The majority of fibrils measured approximately 10 nm diameter and with indeterminate length. Fewer fibrils showed a twist (Figure 1a). AFM images showed fibres with a height of 8-9 nm and a predominant morphology with a smooth cylindrical appearance. However, a minor polymorphic sub-population was observed with a clear helical twist (Figure 1a, b). Thus, prolonged incubation of the fibrils results in a competition between polymorphic structures with a major dominant form of highly ordered, apparent smooth surface fibrils.

**Figure 1.**
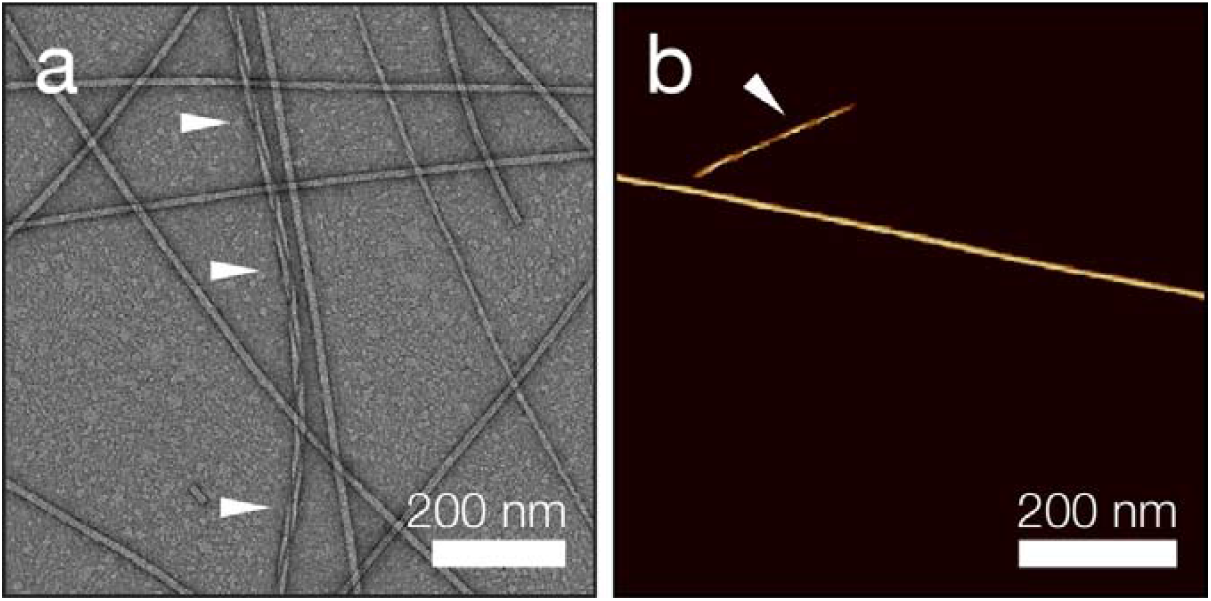
Micrographs showing amyloid fibrils formed by VQIVYK following prolonged incubation. a) TEM shows a major population of smooth fibrils and fewer with a pronounced twist. b) AFM shows very long smooth fibrils and a smaller population of twisted fibrils. White arrows highlight twisted fibrils.

### X-ray fibre diffraction reveals semi-crystalline order within VQIVYK protofilaments

X-ray fibre diffraction patterns were collected from partially aligned VQIVYK fibrils and showed a classical cross-β diffraction pattern with expected 4.86 Å and 10.5 Å diffraction signals perpendicular to one another on the meridian and equator, respectively, arising from hydrogen bonded β-strands, and from the β-sheet spacing, the chain length and the protofilament packing [26] (Figure 2). Remarkably, the diffraction pattern showed that the fibrils within the fibre sample were extremely well oriented and semi-crystalline which gave rise to a far more detailed diffraction pattern than usual for amyloid fibrils (Figure 2, Table 1). Diffraction signals observed at 8.3 Å and 10.5Å show a split appearance as off-equatorial or rho lines (Figure 2, Figure S1).

**Figure 2.**
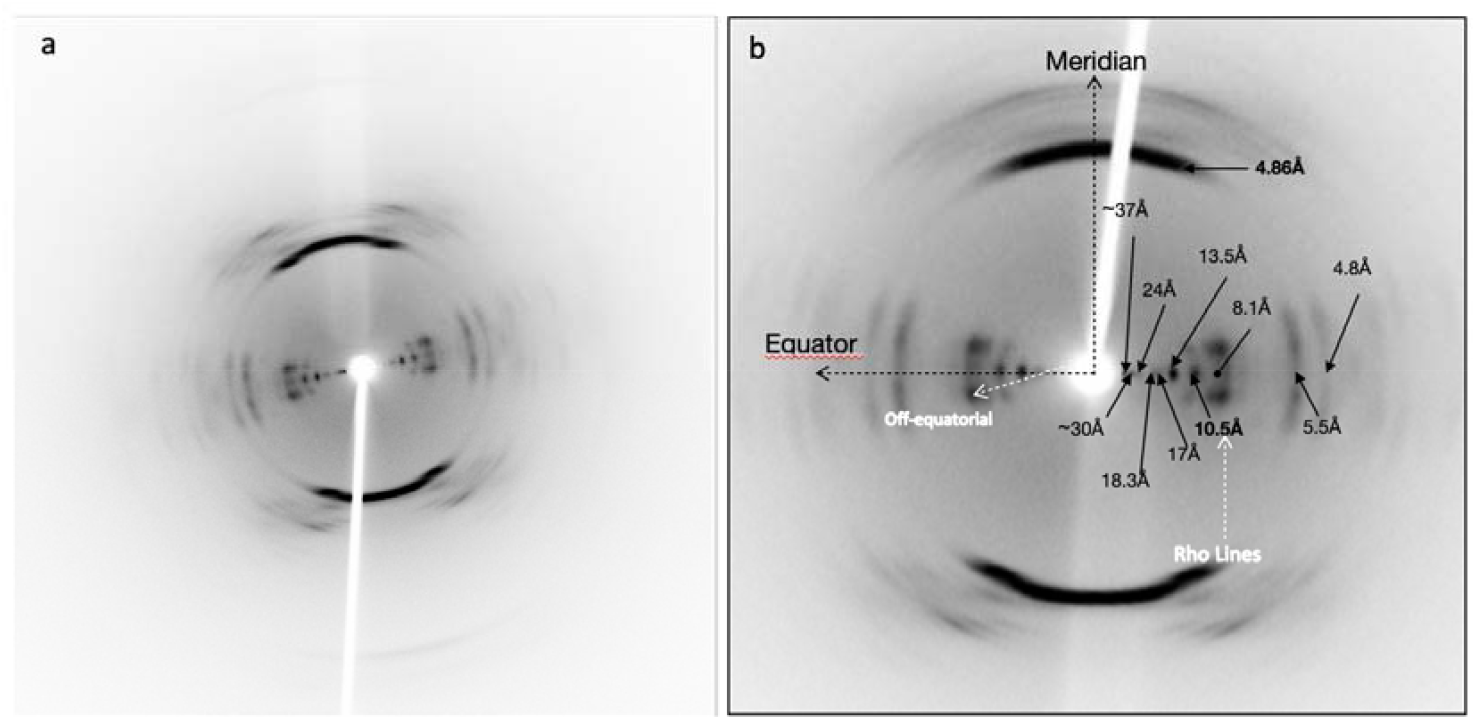
X-ray fibre diffraction from amyloid fibrils formed by VQIVYK. a) Diffraction pattern taken with a distance of 50 mm. b) Diffraction pattern taken at 100 mm specimen to detector distance. Diffraction signals are labelled and the characteristic cross-β signals on the meridian (4.76Å) and equator (10.5 Å) are highlighted in bold. Additional equatorial signals are observed at 10.5Å, 5.5Å and 4.8Å, with lower angle signals labelled. From the equatorial signal at 10.5Å a low intensity-split pattern arises, while 8.1Å displays a split pattern with higher intensity than the signal at the equator. The equator and meridian are labelled with black dotted lines and the off-equatorial and rho line directions are shown in white dotted lines.

**Table 1.** **Diffraction signals position from experimental fibre diffraction pattern from VQIVYK compared to fibre diffraction signals calculated from 2ON9.pdb and twisted model structure in cell** *a*=4.86Å, *b*=61.93Å, *c*=15.41Å; α=90.00º, β=98.11º, γ= 90.00º

| Experimental diffraction signals from VQIVYK fibrils (Å) | Calculated signals from 2ON9.pdb | Calculated from model |
| --- | --- | --- |
| Equatorials |  |  |
| ~37 |  |  |
| ~30 | 30.94 | 30 |
| 24 |  |  |
| 18.3 | 18.52 | 18.52 |
| 17 |  |  |
|  | 16.2 |  |
| 13.5 | 14.1 | 13.11 |
| 10.5 | 10.74 | 10.22 |
| 8.1 | 8.54 | 8.54 |
|  | 6.15 | 6.15 |
| 5.5 | 5.44 | 5.61 |
| 4.8 | 4.99 | 4.9 |
| Meridionals |  |  |
| 4.8 | 4.8 | 4.8 |

The X-ray crystallographical structure of VQIVYK peptide has been solved from microcrystals [19] with unit cell dimensions determined as *a*=4.86Å, *b*=61.93Å, *c*=15.41Å; α=90.00º, β=98.11º, γ= 90.00º which contains two peptides arranged with the same side chains facing each other in a “face-to-face” configuration (2ON9.pdb) (Figure 3a, b). An X-ray fibre diffraction pattern was calculated using the crystal structure coordinates and unit cell dimensions submitted with 2ON9.pdb and the calculated pattern showed a good match with the experimental diffraction pattern from VQIVYK fibrils (Figure 3) (table 1). However, the structure solved from 3D microcrystals necessarily does not incorporate a twist in the β-strands and the calculated pattern does not give rise to observed split in the equatorial signals implying that some detail is missing in the model structure.

**Figure 3.**
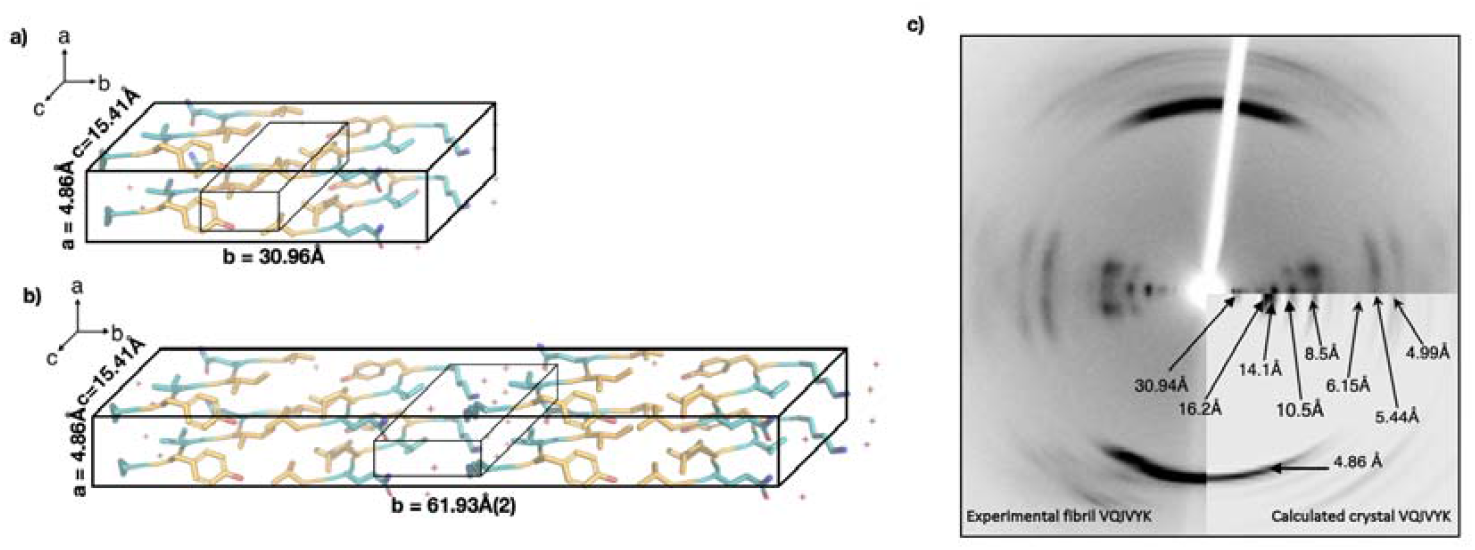
The structure of fibrillar VQIVYK. a) 2ON9.pdb structure is shown in the unit cell derived from the crystal structure [19]. Peptide association depicts the inter-sheet hydrogen bonds between side chains (shown in orange) in a dry steric zipper, highlighted by the smaller box. b) Full unit cell with four peptides along the **b** lattice vector. Red crosses represent the water molecules, that H-bond with blue side chains in the wet steric zipper. Figure was prepared using Pymol [27]. c) Experimental XRDP from fibrillar VQIVYK compared with the pattern calculated from 2ON9.pdb 3D model using the unit cell derived from the crystal structure 2ON9.pdb (a=4.86Å, b=61.93Å, c=15.41Å; α=90.00º, β=98.11º, γ= 90.00º) [19].

Measurement of the angle of the split reflections from the equator is 27.70 ° (Figure S1) and this twist was incorporated into the model structure as a twist of 13.85° for each of the strands within each sheet resulting in a helical pitch of approximately 124 Å for 26 stacked peptides in a highly twisted filament (Figure 3). Calculation of the diffraction pattern from this twisted model structure gives rise to a good match between the calculated and experimental diffraction data (Table 1), and helps to validate the model.

The unit cell from the VQIVYK crystal was further optimised to accommodate the twist and have rise to *a*=4.86Å, *b*≈58Å(2), *c*≈18Å; α=90.00º, β=98.11º, γ= 90.00º. The twisting architecture results in new interactions in the hydrogen bonding of the inter-sheet plane (Fig 4). The side chains from upper layers bond with lower layers, forming a very stable conformation. As a result, Tyr^310^ is closer to the adjacent β-strand than in the planar conformation of the crystal, and aromatic residues make a strong contribution to steric zipper stability [14].

**Figure 4.**
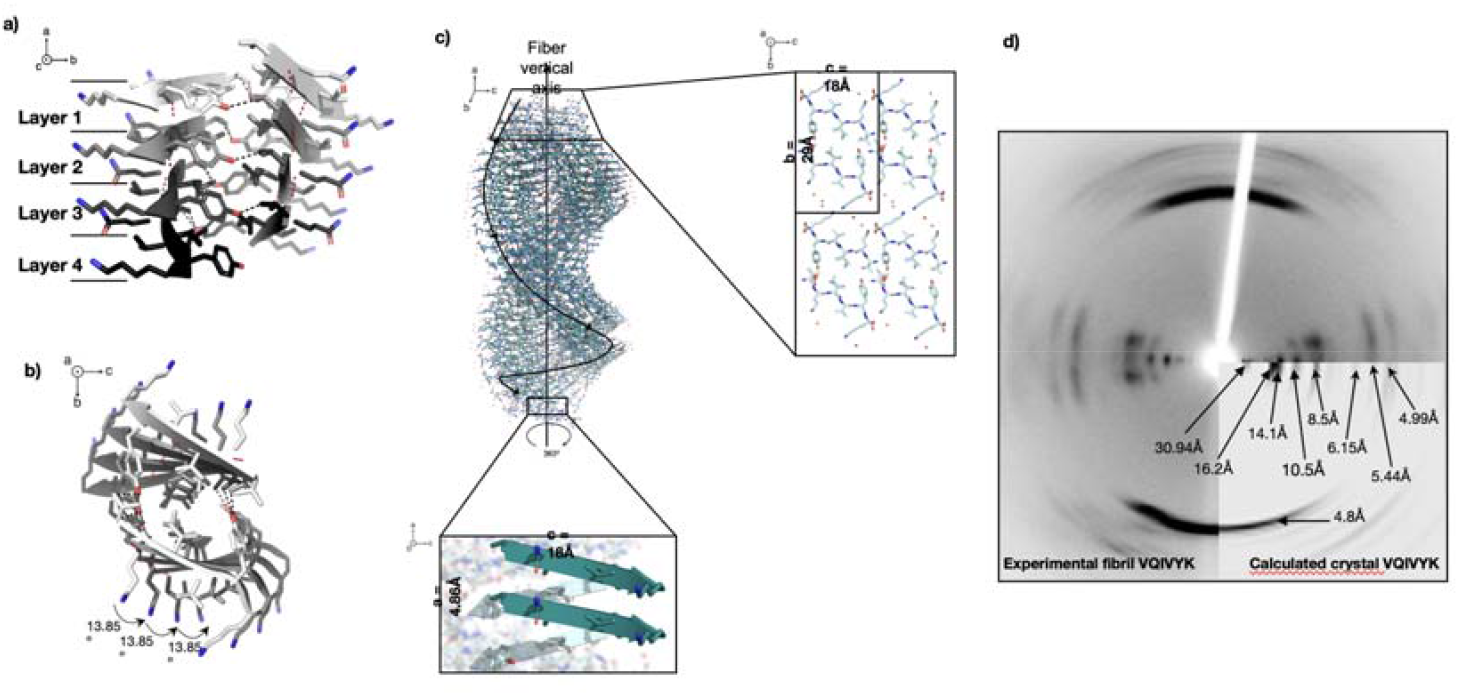
Molecular model of VQIVYK fibrils. **a**) The twist between parallel β-strands allows hydrogen bonding (red dash line) in the lattice vector **a** (4.86Å) forming β-sheets. In the horizontal axes, interactions link side chains from upper layers with lower ones, forming a very stable hydrogen bonding (black dash line) in **b** lattice vector and maintains the intra-sheet bonding distance of 4.86Å. The layers interact via inter-bonding between layers 1—2, 2—3 and 3—4. b) Perspective parallel to the fibre axis (**a** lattice vector) showing the angle applied to the equator of the protofilament. c) The 3D molecular structure of the protofilament shows the twist of 13.85º from layer to layer. The rotation is applied in the equator, resulting in 26 monomers/repeat. d) shows the comparison between calculated pattern from the model created using 2ON9.pdb with an additional 27° twist with experimental data. The optimised model provides an additional diffraction signal that matches with the reflection at 18Å, and split off-meridians that simulate the experimental pattern.

### Structural analysis of individual fibrils by atomic force microscopy shows a mixture of fibril structures within VQIVYK fibril population

X-ray fibre diffraction provides short range information on the core structures of protofilament architecture, while electron microscopy revealed their polymorphic supramolecular organisation. AFM analysis was subsequently employed investigate the polymorph distribution by individual filament structural analysis. Tracing of 59 individual fibrils from AFM images revealed a major fibril population with an apparent smooth cylindrical envelope. This is consistent with highly twisted fibrils where the helical pitch is in the order of 12 nm suggested by the XRFD data. Smooth fibrils have a consistent mean height in the range of 8.6 nm to 9.6 nm (Figure 5), suggesting that the fibrils are formed from multiple protofilaments with ordered core structure seen in the XRFD analysis. Sub-populations of observably twisted fibrils with a clearer pattern of peaks and troughs across the central line height profile compared with the smooth fibrils was also observed with a comparatively larger range of mean height values below that of the major population between 5.7 nm and 9.2 nm, and a large degree of variation in twist and morphology. These sub-populations of fibrils may represent species that were not fully assembled compared with the major population of mature fibrils. The handedness of the helical twist of all fibrils in this population were left-handed, inferring that the handedness for the major population of smooth cylindrical fibrils are likely to be left-hand twisted.

**Figure 5.**
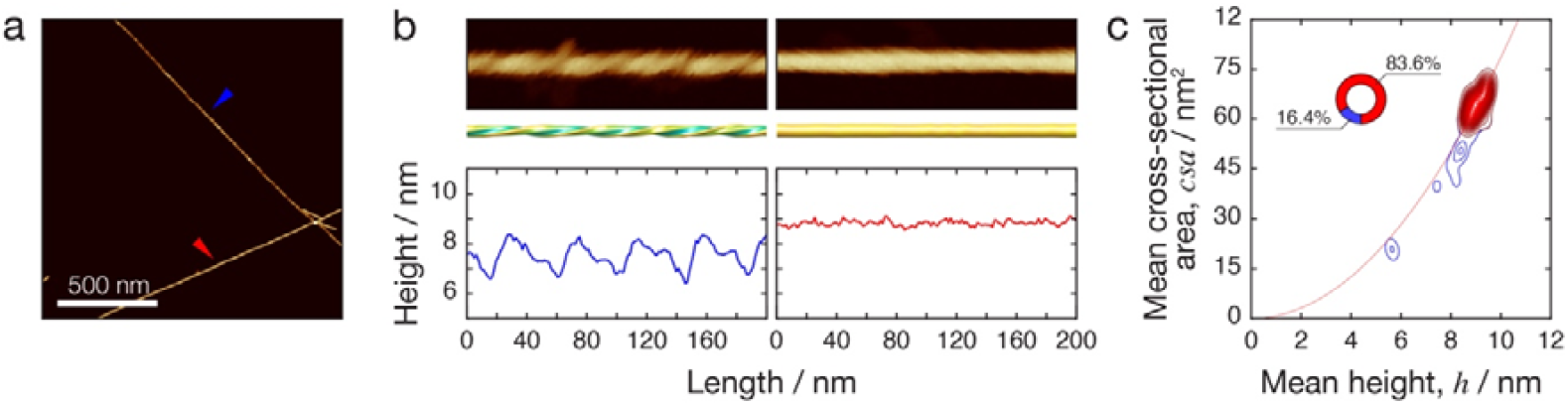
AFM analysis of VQIVYK fibrils shows a major morphology of smooth and cylindrical fibrils and minor sub-population of fibrils with various twisting morphologies. A) AFM image of VQIVYK fibrils. Scale bar represents 500 nm. B) Analysis of two fibrils with different morphologies. The fibrils shown in b) are those highlighted with blue and red arrows in a), respectively. Top panels show straightened AFM images of a 200 nm section of the fibrils. The middle panels show reconstructed 3D envelopes of the fibrils. The bottom panels show the height profiles of the fibrils across the central lines of the straightened filaments. A 200 nm segment along the length of the reconstructed fibrils is shown for comparison. C) Mean height of the fibril central line plotted against the mean cross-sectional area estimated by 3D contact point reconstruction AFM. Distribution densities from 61 individual fibrils are plotted. The smooth fibrils are shown in red and fibrils with clear peak-and-trough pattern are shown in blue. Inset shows the relative size of the two sub-populations. A black line shows values expected for perfectly cylindrical fibrils.

### Integrating structural information from XRFD and AFM allowed creation of multi-scale structural models of VQIVYK fibrils

Reconstruction of three-dimensional envelopes from traced and straightened fibril height data enabled the average cross-sectional area to be calculated for each individual fibril. Thus, it is possible to estimate the number of pairs of VQIVYK β-sheets that pack laterally to form the fibril ultrastructure. Simple comparison of areas between the half-unit cell of paired β-sheets and cross-sections of the smooth fibrils could indicate that between 11-14 peptide pairs could be packed to form smooth fibrils’ cross-sections (Figure 6). A different arrangement with a more elongated cross-sectional shape would be expected for the minor twisted fibril populations and intrafibrillar variation in the number of peptide pairs or their arrangement is suggested from the morphology of the fibrils as shown in Figure 6.

**Figure 6.**
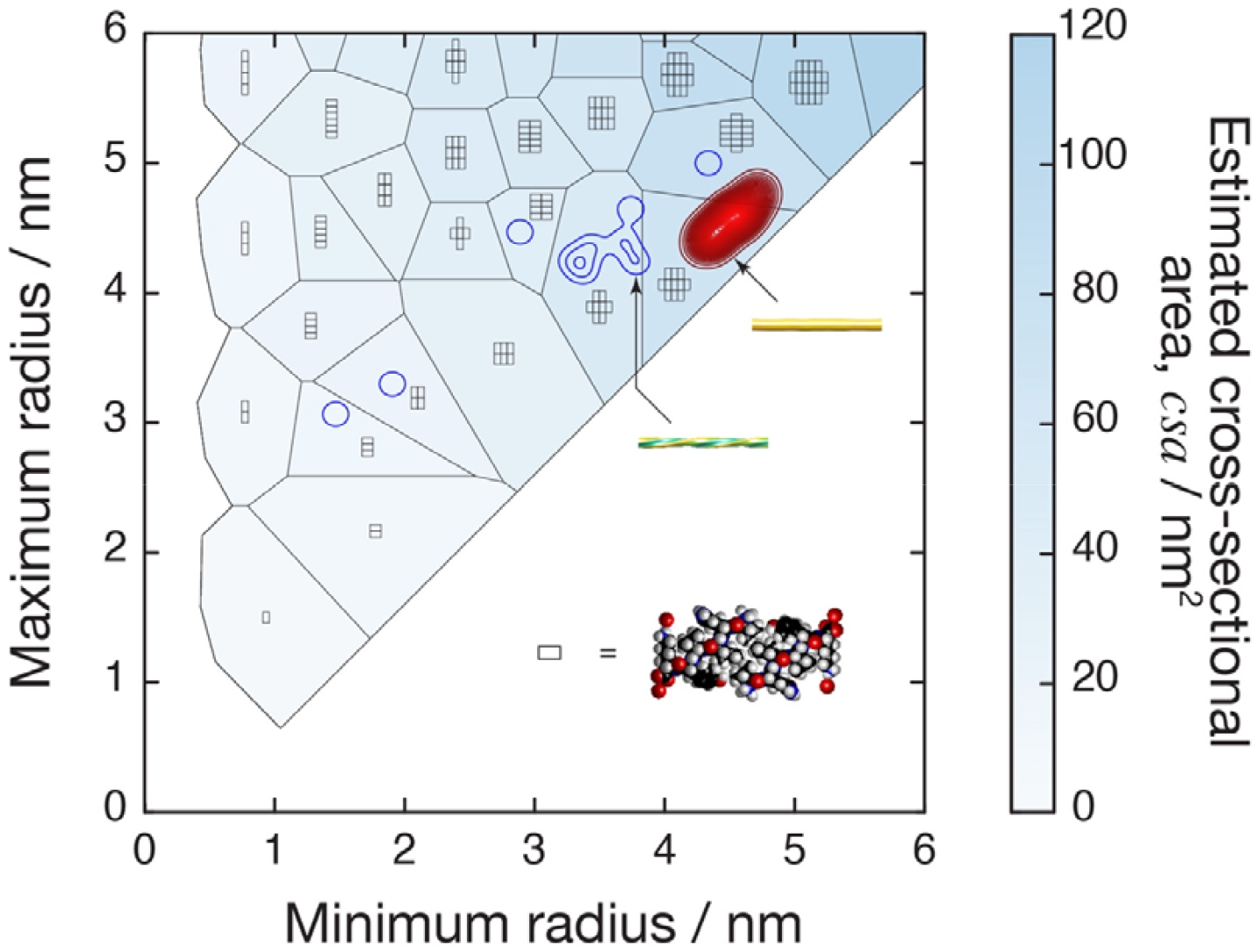
Possible packing arrangements of VQIVYK unit cell cross-sections. Arrangements of pairs of VQIVYK β-sheets in a half-unit cell modelled from the XRFD (symbolised by the rectangles) mapped onto the cross-sectional dimensions of fibril envelopes reconstructed from the AFM data. Possible packing arrangements tested are those with fully convex cross-sections and two-fold symmetry. Regions consistent with each of the arrangements are shown with an illustration of the packing of the peptide pairs (rectangles), and the colour of the regions indicate cross-sectional area estimated by the cross-sections’ convex hull area. This is mapped onto the polymorph distribution of the fibrils visualised by their minimum radius plotted against the maximum radius obtained by 3D CPR-AFM. The distribution of smooth fibrils are shown in red and the distribution of fibrils with clear peak-and-trough pattern are shown in blue.

## Conclusions

Understanding the architecture of amyloid fibrils has made major advances since our understanding of the generic cross-β structure [1, 28] with microcrystallography and cryo-electron microscopy both providing exciting details regarding the interactions between side chains in steric zippers and the organisation of in-register parallel stacking of individual protein chains. However, the understanding of how the β-sheets associate and build to create twisted mature fibrils remains unclear. Here, using a powerful combination of X-ray fibre diffraction to define the molecular architecture of the protofilament cores in the sample, and atomic force microscopy to provide a details molecular envelope for each individual filament, we have gained valuable insights into the overall organisation of peptides within an amyloid fibril and concluded that the fibrils are assembled from protofilament building blocks with highly ordered and common cores. We show that a fine twist within the sheets creates a tight helix which results in very ordered fibril cores. The atomic force microscopy reinforces the structural model and gives further insight into how protofilaments associate to result in structural polymorphism. Furthermore, we show, from this prolonged incubation of an amyloidogenic peptide, that this can converge to form extremely stable, ordered fibril core structures.

## Acknowledgements

YA wish to thank ministry of higher education and scientific research of Iraq for the support and fund. This work was supported by funding from the Biotechnology and Biological Sciences Research Council (BBSRC), UK grant BB/S003657/1, BB/S003312/1 and BB/Z516880/1 and Engineering and Physical Sciences Research Council (EPSRC), UK DTP grant EP/R513246/1 (LL).

## Open research

## Supplementary information

### Materials and Methods

Figure S1. Close up view of equatorial region of the diffraction pattern.

