## Supplementary material for "The architecture of amyloid fibrils formed by a human tau-derived hexapeptide VQIVYK": SI

*Equal authors

^1^ Sussex Neuroscience, School of Life Sciences, University of Sussex, Falmer, E Sussex, BN1 9QG

^2^ Chemistry Department, College of Science, Mustansiriyah University, Baghdad, Iraq

3. School of Biosciences, University of Kent, Canterbury CT2 7NJ, UK

**Materials and Methods**

**Amyloid fibril formation**

Capped VQIVYK was purchased from JPT at 97% purity as a TCA salt. The peptide was dissolved in Milli-Q 0.22 μm filtered water at 1 mg/ml and was allowed to self-assemble at room temperature for approximately two years.

**Transmission electron microscopy**

4 μL of fibril solution was applied to formvar coated 400 mesh copper grids (Agar Scientific) and allowed to adhere for 60 seconds before blotting, washing with 4 μL of filtered Milli-Q distilled water and then incubating with 0.22 μm filtered uranyl acetate (2% w/v) for 30 seconds and then washed twice with 4 μL of filtered Milli-Q distilled water and allowed to air dried. The grids were examined using a JEOL JEM1400-Plus Transmission Electron Microscope (TEM) operated at 120 kV and images were collected at 20-40K magnification using a Gatan OneView camera (4Kx4K). Images were recorded at 25 fps with drift correction using Digital Micrograph (GMS3, Gatan).

**Atomic force microscopy data collection and analysis**

The pre-assembled fibril sample was diluted 1/50 in sterile filtered Milli-Q water adjusted to pH 2.0 using a dilute solution of HCl. Immediately after dilution, 20 μl samples were deposited onto freshly cleaved mica surfaces (Agar scientific, F7013) and incubated for 10 min. Following incubation, the sample was washed with 1 ml of the filter sterilised HCl solution and dried using a stream of nitrogen gas. Fibrils were imaged using a Multimode 8 AFM with a Nanoscope V (Bruker) controller operating under peak-force tapping in the ScanAsyst mode using Bruker ScanAsyst probes (silicon nitride triangular tip with nominal tip radius of 2 nm and nominal spring constant of 0.4 N/m, Bruker). Height channel images, with scan sizes of 3 × 3 μm, 5 × 5 μm, or 8 × 8 μm were collected with 2048 × 2048 pixels each. Nanoscope analysis software (Version 1.5, Bruker) was used to process the image data by flattening the height topology maps to remove tilt and scanner bow. Flattened image data was imported into Trace_y software ^[1]^ where all subsequent analysis was carried out. Fibrils were traced and digitally straightened and the height profile for each fibril was extracted from the centre contour line of the straightened fibrils. Straightened fibril traces were corrected for the tip-convolution effect using an algorithm based on geometric modelling of the tip-fibril contact points, followed by 3D reconstruction of the fibril envelopes as previously described^[1]^. The method involves determining the helical pitch of each fibril through Fourier transform of the fibril centre contour heights. Where this was not possible due to lack of any clear periodicity in the central line height profile of the fibril, the mean pitch of fibrils with detectable pitch was used. It was previously confirmed on three smooth fibrils that while varying the pitch from 10 to 100 pixels the mean difference between maximum and minimum of mean cross-sectional area values is 0.23 nm^2^ indicating that using an average pitch in the case of these smooth fibrils has comparatively little effect on the cross-sectional area measurements from the reconstructed fibril envelope models.

**X-ray fibre diffraction data collection and analysis**

10 μL droplet was placed between two wax-tipped capillary tubes arranged in a petri dish at 4 °C and allowed to dry for several days. The resulting bundle of fibrils on a capillary tube was mounted on a goniometer head and X-ray fibre diffraction data was collected using a Rigaku CuKα rotating anode and RAxis 4++ detector. Exposure times were 30s or 60s with a specimen to detector distance of 50 or 100 mm respectively. Patterns were converted to tiff format and inspected using CLEARER ^[2]^. The patterns were *centred* and radially averaged, and the diffraction signal positions were measured using the *peak finder* tool in CLEARER. The signals positions were further confirmed using *Zoom and Measure* tool. Values that diverged from the equatorial were measured using the angle in pixels from the origin and calculated using trigonometry. Unit cell dimensions were derived using *unit cell determination* tool following the input of equatorial diffraction signals.

**Modelling**

Models were prepared using the X-ray crystallography structure 2ON9.pdb^[3]^ as a starting model. A model structure was generated using a python script that imposes specific translation and rotation parameters and these models were generated using Pymol ^[4]^.

**Diffraction pattern calculation**

Diffraction patterns were calculated from PDB coordinates generated from Pymol ^[4]^ using the *Diffraction simulation tool* in CLEARER and the determined unit cell parameters^[2]^.

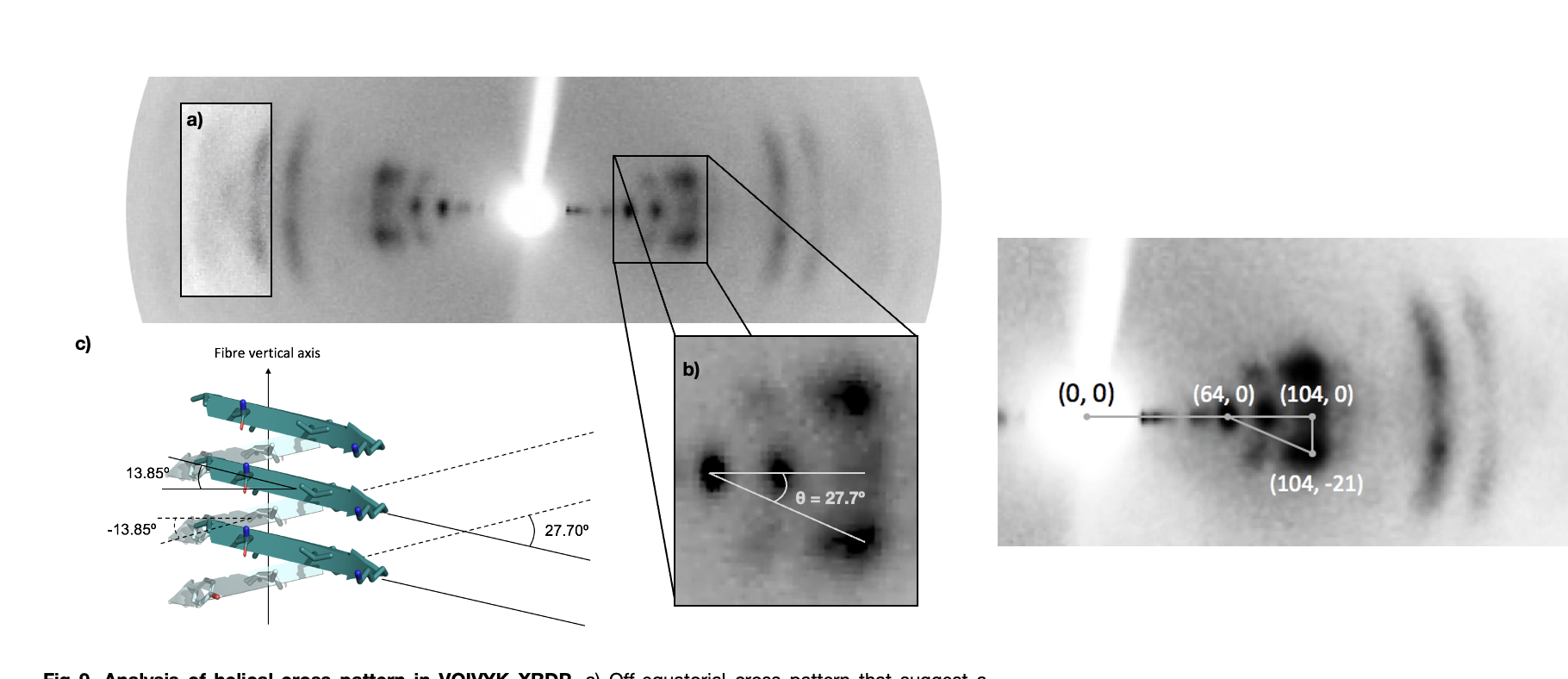

Figure S1. X-ray fibre diffraction pattern analysis of fibrils formed by VQIVYK. a) highlights the high angle split reflections. b) shows the angle between equatorial and the off-equatorial signals. c) shows the organisation of the peptides to accommodate the twist.
